# Accelerating Drug Discovery and Repurposing by Combining Transcriptional Signature Connectivity with Docking

**DOI:** 10.1101/2020.11.25.399238

**Authors:** Alexander W. Thorman, James Reigle, Somchai Chutipongtanate, Behrouz Shamsaei, Marcin Pilarczyk, Mehdi Fazel-Najafabadi, Rafal Adamczak, Michal Kouril, Ardythe L. Morrow, Maria F. Czyzyk-Krzeska, Robert McCullumsmith, William Seibel, Nicolas Nassar, Yi Zheng, David Hildeman, Andrew B. Herr, Mario Medvedovic, Jarek Meller

## Abstract

The development of targeted treatment options for precision medicine is hampered by a slow and costly process of drug screening. While small molecule docking simulations are often applied in conjunction with cheminformatic methods to reduce the number of candidate molecules to be tested experimentally, the current approaches suffer from high false positive rates and are computationally expensive. Here, we present a novel *in silico* approach for drug discovery and repurposing, dubbed *connectivity enhanced* Structure Activity Relationship (*ce*SAR) that improves on current methods by combining docking and virtual screening approaches with pharmacogenomics and transcriptional signature connectivity analysis. *ce*SAR builds on the landmark LINCS library of transcriptional signatures of over 20,000 drug-like molecules and ~5,000 gene knock-downs (KDs) to connect small molecules and their potential targets. For a set of candidate molecules and specific target gene, candidate molecules are first ranked by chemical similarity to their ‘concordant’ LINCS analogs that share signature similarity with a knock-down of the target gene. An efficient method for chemical similarity search, optimized for sparse binary fingerprints of chemical moieties, is used to enable fast searches for large libraries of small molecules. A small subset of candidate compounds identified in the first step is then re-scored by combining signature connectivity with docking simulations. On a set of 20 DUD-E benchmark targets with LINCS KDs, the consensus approach reduces significantly false positive rates, improving the median precision 3-fold over docking methods at the extreme library reduction. We conclude that signature connectivity and docking provide complementary signals, offering an avenue to improve the accuracy of virtual screening while reducing run times by multiple orders of magnitude.

## Introduction

Accelerating the pace of drug discovery and repurposing is paramount for the development of treatments for rare diseases or personalized treatment options for precision medicine, and the ability to respond to public health crises, such as the COVID-19 pandemic. Systematic efforts for drug discovery have used high throughput *in vitro* or *ex vivo* screening approaches, often in conjunction with an initial *in silico* screening of small molecule libraries. These efforts have resulted in a large number of candidate compounds targeting the druggable part of the genome^1–3^. Parallel advances in pharmacogenomics and large-scale candidate drug profiling in cell lines and other model systems, such as Connectivity Map^4^, NCI60^5^ and Cancer Cell Line Encyclopedia^6^, or GDSC^7,8^, have further revolutionized drug discovery, target and mode of action prediction, and repurposing. For example, transcriptional signature connectivity analysis has been used to identify drugs that may reverse a signature of a disease state or that may have the same mode of action because of the similarity of their signatures^4,9–11^.

The LINCS consortium has recently compiled a library of transcriptional signatures for over 40,000 drug-like molecules as well as over 6,000 gene knockdown (KD) and overexpression constructs in multiple cell lines^12,13^. As a result, LINCS transcriptional signatures can be used to directly correlate downstream transcriptional responses induced by chemical perturbations with those induced by loss or gain of function of the target protein. By enabling the direct exploration of *drug-gene* relationships on a previously unattainable scale, using significant subsets of both: the drug-like universe of small molecules and druggable genome, LINCS provides a unique big data resource for pharmacogenomics^13–15^ that is explored here with the goal of identifying candidate inhibitors of a specific protein target.

It should be emphasized that similar downstream transcriptional signatures may result from the loss of function of multiple upstream proteins in signaling cascades or pathways converging on the same transcriptional targets. Of particular relevance are signaling cascades involving multiple kinases and phosphorylation events between a growth receptor and a transcription factor in many types of cancer^16^. Thus, the analysis of concordance between signatures of small molecules and the target gene knock-down can identify candidate molecules that effectively lead to the loss of function as pathway inhibitors, and not necessarily a specific target inhibitor, as illustrated in Figure 1. The left panel in the figure pictorially represents a signature connectivity analysis to identify putative inhibitors of SRC by searching for candidate small molecules whose signatures are concordant, i.e., positively correlated with the SRC KD signature. Note that all 3 compounds targeting the EGFR – SRC – JUN signaling cascade are ‘concordant’, although only one of them targets SRC directly.

**Figure 1:**
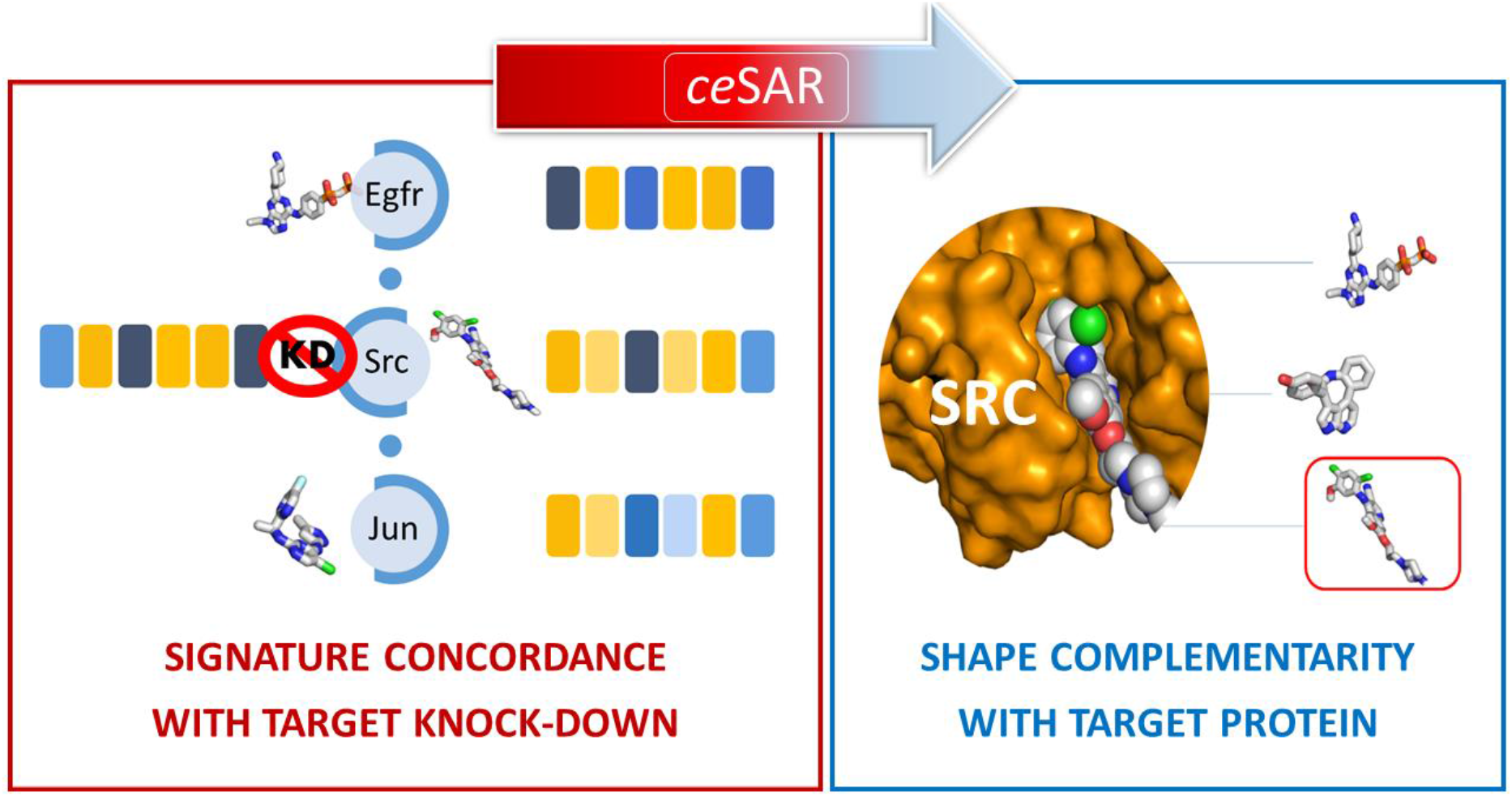
The overall principle of the new *connectivity enhanced* Structure Activity Relation (*ce*SAR) approach that ranks candidate molecules by their similarity to LINCS analogs with signatures concordant to those of the target gene KDs (left panel), and can be subsequently combined with docking simulations to assess the shape complementarity with specific protein targets (right panel). Transcriptional signatures are defined as down-and up-regulated genes with the corresponding differential expression values, represented as blue and yellow boxes in the left panel, respectively. The fictitious SRC KD signature consists of 6 genes, with genes 1, 3, and 6 down-regulated and genes 2, 4, and 5 up-regulated. All 3 compounds targeting the EGFR – SRC – JUN cascade result in signatures concordant with that of SRC KD, but only the actual SRC inhibitor fits the binding pocket in docking.

To achieve the specificity to a target in a pathway, the predicted binding affinity to a target protein may be used to complement the signature connectivity-based approach. This is illustrated in the right panel in Figure 1 where the actual SRC inhibitor is shown as predicted to have the lowest binding energy, and thus selected as consensus candidate. In this context, various *in silico* docking techniques have been widely used to computationally predict binding affinities between small molecules and their (structurally resolved) targets, often coupled with Structure-Activity Relationship (SAR) analysis of chemical analogs for top ranking candidate molecules^17,18^.

Here, we present a novel approach for accelerating drug discovery and repurposing, dubbed *connectivity enhanced* Structure Activity Relationship (*ce*SAR), that combines these two principles. Capitalizing on the LINCS library of transcriptional signatures (denoted as L), *ce*SAR combines drug and target transcriptional signature connectivity analysis with efficient chemical similarity search and virtual screening approaches. For a gene target and a library of candidate compounds to be screened, a subset of ‘concordant’ LINCS small molecules is first identified to include only those compounds that have signatures concordant with a target gene knock-down (or over-expression) signature. The library of candidate compounds is then reduced by using a fast chemical similarity search, optimized for sparse binary fingerprints of chemical moieties, to identify those compounds that are, with some Jaccard similarity^19^ threshold, structural analogs to a LINCS molecule with transcriptional concordance to the genetic knock-down (or overexpression). The resulting small subset of compounds can be subsequently re-scored in conjunction with docking, using a consensus ranking to filter out likely pathway, but not target protein inhibitors.

## Results

In order to assess the new method and test the hypothesis that combining the principles of signature connectivity and shape complementarity can improve drug discovery by reducing the level of false positives and reducing computational cost of virtual screening, we systematically evaluate the performance of *ce*SAR and compare it with the results of *Autodock^20^* and *MTiOpenScreen*^20^, using a subset of targets from the DUD-E benchmark, which is widely used in the assessment of docking and virtual screening methods^22^.

### Candidate molecule ranking using ceSAR

For a library of small molecules, *Q*, and a target gene t with at least one consensus shRNA knock-down transcriptional signature available in LINCS, *t* ∈ *L, ce*SAR ranks candidate compounds by identifying their closest chemical analogs in the LINCS library of transcriptionally profiled chemical perturbagens, *k* ∈ *L*, that result in signatures concordant to those of the target KDs. For each *q* ∈ *Q*, the following similarity score is computed as a basis for ranking:

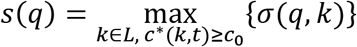

where *σ*(*q,k*) is the Tanimoto coefficient (Jaccard similarity measure)^19^ between compounds *q* and *k* represented as binary fingerprints, and *c*\*(*k,t*) is the maximum concordance (over all cell lines for *t*, and cell line, concentration, exposure time tuples for *k)* between the signatures of chemical perturbagen *k* and genetic knock-downs of *t. i*LINCS correlation-based concordance measure is used here^23^, and the threshold for significant concordance is set to *c*_0_ = 0.2, as discussed in the Methods sections.

Note that the similarity score, *s*(*q*), is in fact the Tanimoto coefficient for the closest ‘concordant’ LINCS analog of *q*, and thus is a real number between 0 and 1. By increasing the similarity threshold, *s*_0_ ∈ [0,1], one can reduce the initial library to a (hopefully enriched into true positives) subset that can be used for further analysis and validation by taking only those compounds *q* that receive a score larger than *s*_0_.

While different forms of combining chemical similarity and concordance measures into a composite score can be considered to potentially improve the performance, such as machine learning-based ensemble consensus classifiers discussed in *Supplementary Materials*, we deliberately use here this very simple form of the method, which will be referred to as S throughout the manuscript, to evaluate the advantages of *ce*SAR.

### Fast exact chemical similarity search

The LINCS library of drug-like molecules comprises over 40,000 compounds, while the user defined library of small molecules *Q* to be ranked and reduced by identifying ‘concordant’ LINCS analogs, can be quite large in the context of virtual screening. An efficient solution for computing the Jaccard similarity measure (Tanimoto coefficient) and retrieving the closest matches for the case of sparse binary fingerprints is used here to address this computational bottleneck and accelerate the *ce*SAR search. As shown in the Methods section, by using pre-processing of the reference data set of compounds (here LINCS library), the computation of similarity scores between a query compound *q* and database compounds can be limited to only those columns in the binary fingerprint where *q* is in the minority state, which is assumed to be 1, while also optimally exploiting the sparsity in each column of the fingerprint across database compounds by pre-computing indexes of database compounds in the minority state at that column. The resulting algorithm, dubbed *minSim* (for *minority Sim*), optimally exploits the sparse nature of binary fingerprints commonly used for fast chemical similarity search without using approximate techniques, such as those based on hashing^24–26^. For the retrieval from the LINCS library, *minSim* provides between 60 and 150-fold speed-up for different DUD-E datasets compared with traditional approaches (see Supplemental Table 1).

### Consensus re-ranking using docking to improve ceSAR

The initial *ce*SAR search, as defined above, can be subsequently combined with docking simulations to achieve higher specificity for a target at hand. For the purpose of systematic benchmarking and assessment of robustness of the new method, we consider several forms of consensus, starting from the entire library and performing the initial *ce*SAR search (S) and docking simulations for all compounds in the library to derive the consensus ranking, referred to as C_100_, or starting from a subset of the library, first reduced using the signature connectivity based filter S. When the library is first reduced to the top 5% or 1% of the library, the consensus form of *ce*SAR is referred to as C_5_ and C_1_, respectively. Note that C_1_ reduces the computational time 100-fold compared to docking, as only the top 1% of the library identified by using S needs to be screened by docking. While more complex, machine learning-based models to combine signature connectivity related features and docking scores are benchmarked in *Supplementary Materials*, here we present the results of a simple consensus that aims to minimize the risk of overfitting, and defines the combined rank as the geometric average of signature connectivity and docking based ranks.

### Benchmarking of ceSAR

The Directory of Useful Decoys Enhanced (DUD-E) benchmark was developed to determine the success rate of virtual screening methods and assess their ability to discriminate between the known binders and carefully design sets of decoys that are unlikely to bind to target proteins. To evaluate the performance of *ce*SAR, we used a subset of 20 targets from the original DUD-E benchmark^22^ that had gene knockdowns available within LINCS. More details regarding the benchmark datasets used here are provided in the *Supplementary Materials*. The simple signature connectivity-based *ce*SAR search, referred to as S for signature-based (or *Sig2Lead)*, is compared with the traditional virtual screening using *AutoDock* v. 4.2 docking program and the set of original DUD-E crystal structures for target proteins, referred to as A for *Autodock,* and with the combined approach, denoted as C for consensus-based, that uses the simple search (S) first to reduce the library, and then applies docking (A) to such reduced subset to derive the consensus ranking as defined in the previous section.

In addition, a simple baseline (denoted B) that ignores signature concordance while identifying the closest LINCS analogs of DUD-E compounds, and thus simply reflects biases in coverage for inhibitors of different targets in LINCS, is included to assess the contribution of signature concordance. The results are summarized in Figures 2, 4 and 5 in terms of precision curves to quantify the enrichment into true binders upon the reduction of the DUD-E datasets to small subsets amenable to further validation. Note that the precision, or positive predictive value, defined as *PPV* = *TP/(TP* + *FP*), where *TP* denotes the number of true positive predictions and *FP* the number of false positive predictions, captures how many candidate compounds, selected by *in silico* ranking, are in fact true binders. Thus, the precision or *PPV* measures the likelihood of successfully identifying an inhibitor through experimental validation for a small subset of the library reduced here by using *ce*SAR or docking, which arguably is the most relevant measure of success for virtual screening methods.

**Figure 2:**
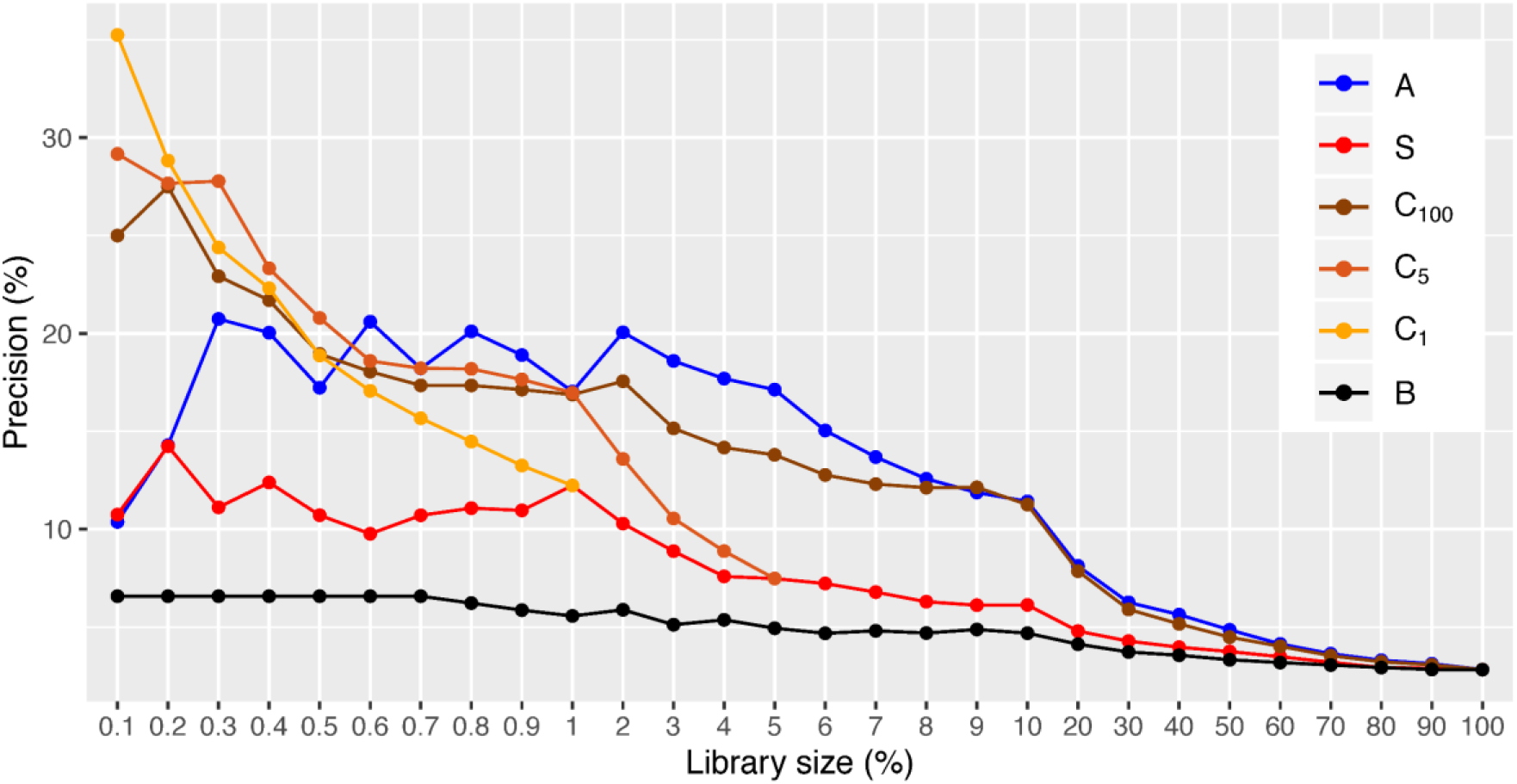
The median precision (or positive predictive value) for 20 DUD-E targets as a function of library size for the simple *ce*SAR search using *Sig2Lead* (S), docking using *AutoDock* (A), and consensus approaches that combine the initial library reduction by *Sig2Lead* with *AutoDock* for the top 1%, 5% and 100% library subsets (C_1_; C_5_ and C_100_, respectively), compared with a simple baseline method (B) that ignores signature connectivity and uses only chemical similarity to LINCS compounds for library reduction.

We would like to emphasize that given the negligible computational cost compared with docking (see Figure 7) and the low barrier to applying *ce*SAR, one might consider a success even modest levels of enrichment or increased precision as the library is reduced. The other measure of success is the level of improvement over the baseline that measures the signal due to signature connectivity. As can be seen from Figure 2, the simple form of *ce*SAR (S) in fact performs significantly better than the baseline (p-value of 6.1 × 10^-10^ using Kolmogorov-Smirnov test), while the consensus form of *ce*SAR significantly outperform both A and S (including C_1_ that yields Kolmogorov-Smirnov p-values of 2.8 × 10^-3^ and 1.1 × 10^-8^ in comparison with A and S, respectively). Importantly, these improvements are obtained for the most reduced, and thus arguably most relevant, library sizes, where the consensus approaches achieve median precision of more than 25% (35% for C_1_), compared with just 10% for docking.

Thus, on the DUD-E benchmark (using the original target conformations and binding sites), docking is successful in eliminating the most unlikely binders by using shape complementarity and the predicted binding energies, leading to initial success and higher precision (and enrichment) at the level of 5 or 10% of the original library. However, docking struggles to correctly rank true positives and the remaining (more challenging) true negatives, resulting in a drop of accuracy as the size of the library is reduced further. Note that DUD-E datasets comprise tens of thousands of molecules, so reduction to less than 1% of the library size is desirable to reduce the number of compounds for testing.

In terms of the distribution of precision values over 20 DUD-E targets, the results of the consensus-based *ce*SAR method (C_1_) and docking (A) are becoming statistically indistinguishable at 0.5% library size, while providing significant speed-ups since only a small fraction (1%) of the library needs to be re-scored using consensus with docking (see Figure 7). On the other hand, the simple *ce*SAR search (S) that can be performed on a personal laptop within minutes, achieves results statistically indistinguishable from docking at 0.1% library size, with both methods yielding a median precision (or positive predictive value) of about 10% at this furthest library reduction (see also Figure 2).

Note that 0.1% library size on average corresponds to only about 20 compounds to be tested. At this furthest library reduction, the fraction of true binders among the candidate compounds selected by using the C_1_ consensus *ce*SAR approach, which combines signature connectivity analysis with docking for the top 1% of the library ranked by S, is equal or greater than 35% for half of DUD-E targets. We conclude that C_1_ consensus method provides the best trade-off between speed and accuracy on the DUD-E benchmark, while the performance of the consensus-based *ce*SAR methods (C_1_, C_5_ and C_100_) is robust with respect to the choice of the top library subset for integration with docking. This is further illustrated by the distribution of top true positive rank for individual DUD-E targets (see Figure 3) that shows good performance of consensus methods. Similar results are obtained on a subset of DUD-E targets (as well as on the original DUD benchmark) using *MTiOpenScreen* docking server (see *Supplementary Materials).*

**Figure 3:**
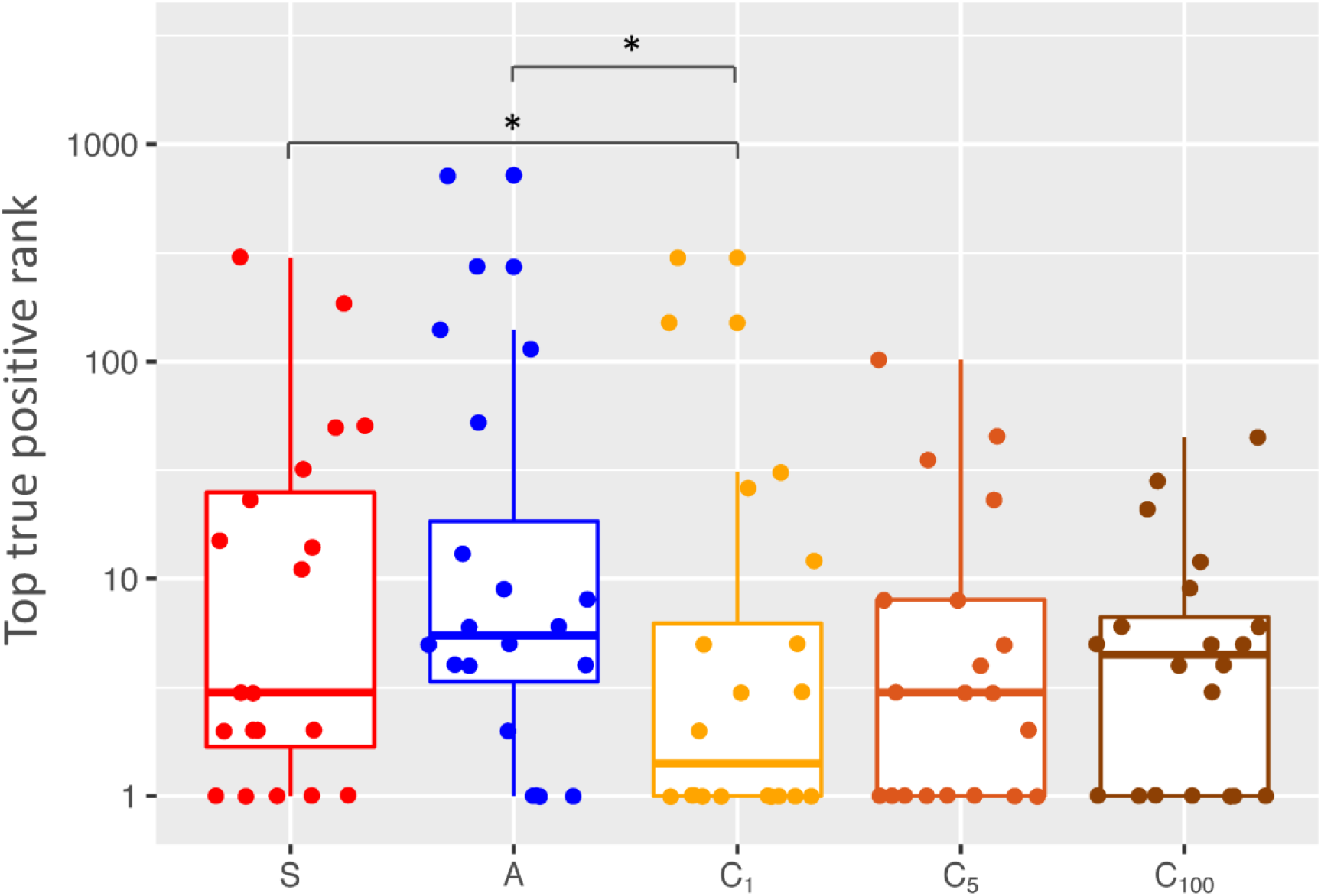
The distribution of the top true positive rank for 20 DUD-E targets at 0.1% library reduction for the simple *ce*SAR search using *Sig2Lead* (S), docking using *AutoDock* (A), and consensus approaches C_1;_ C_5_ and C_100_ (see also Figure 1). Note that C_1_ improves significantly on *Sig2Lead* or *AutoDock* alone with Wilcoxon test p-values < 0.05 (without outliers) indicated by stars, while the other two consensus approaches avoid failures, i.e., targets without true positives in the top 100 candidates.

Importantly, *ce*SAR is more robust compared to docking, which performs very well for some targets while also failing completely for several targets at that library size. This is illustrated in Figures 4 and 5 using precision curves for individual targets and comparison of the area under the precision curve at different library sizes, respectively. As can be also seen from Figure 6, at the most extreme library reduction considered here (0.1% library size), *AutoDock* fails to retain any true positives and thus yields precision of 0% in 8 out of 20 cases, compared with 7 for the simple *ce*SAR search, and only 4 such failures for the C_1_ consensus method. These trends also hold in terms of the number of targets for which none of the true positives is ranked among top 100 candidates (Figure 3).

**Figure 4:**
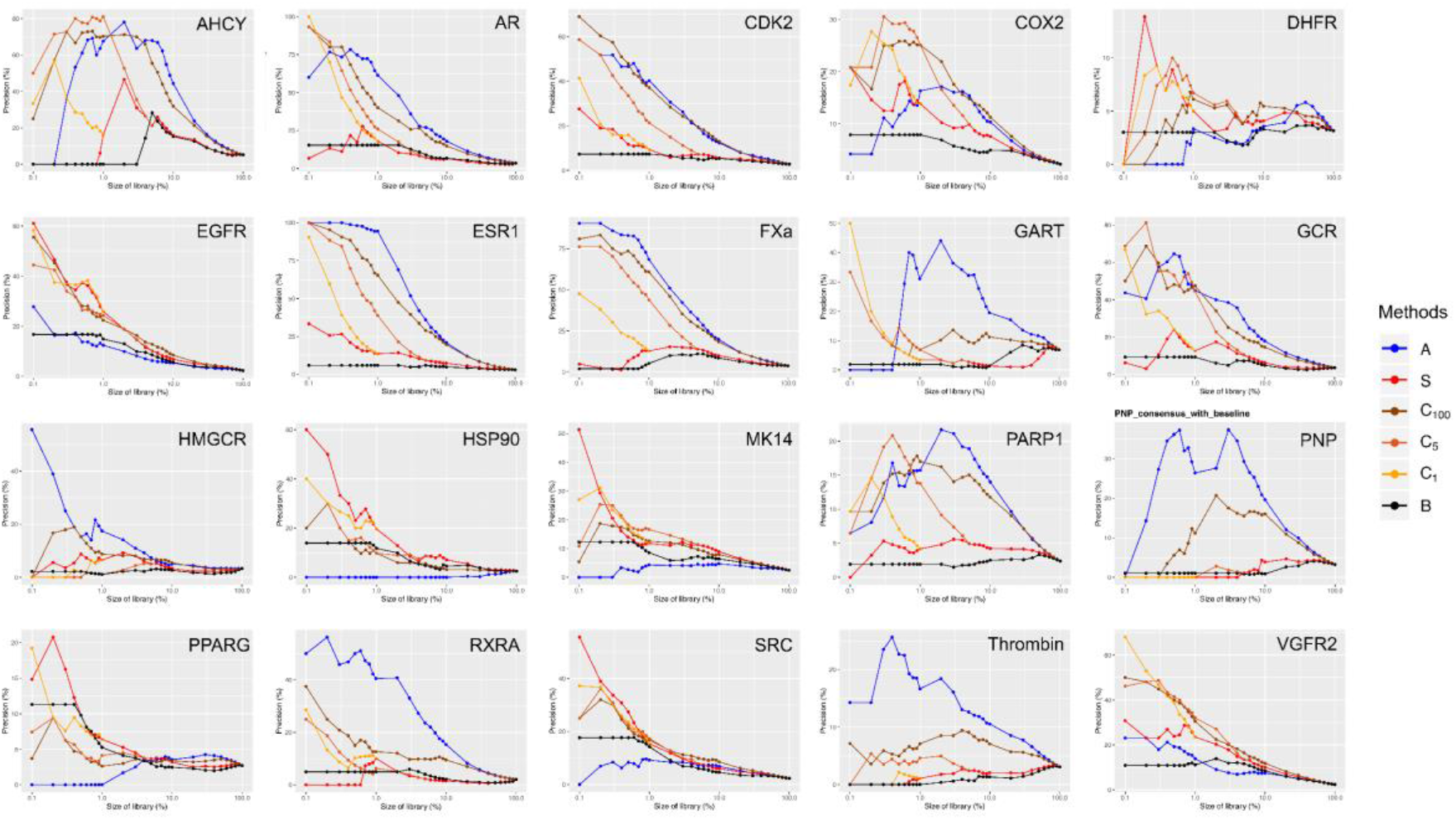
Individual precision curves as a function of library size for DUD-E targets. Either *Sig2Lead* alone (Red, S) or consensus *ce*SAR methods (Yellow, C_1_, Orange, C_5_, Brown, C_100_) improve in precision over *AutoDock* alone (Blue, A) in 12 of the 20 cases, while performing on par with docking for additional 4 targets at the most reduced library sizes.

**Figure 5:**
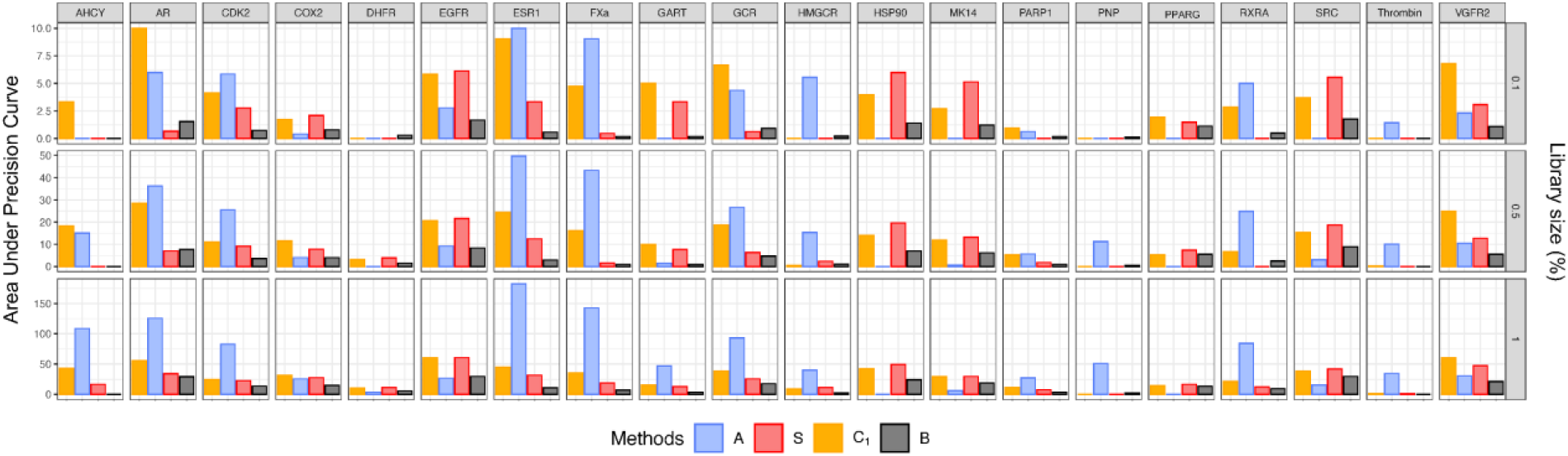
Performance of the consensus *ce*SAR approach (C_1_, Yellow), docking (A, Blue), the simple *ce*SAR search (S, Red), and the baseline method (B, Black) on DUD-E benchmark in terms of the area under the precision curve (arbitrary units for comparison between targets) at different library size.

**Figure 6:**
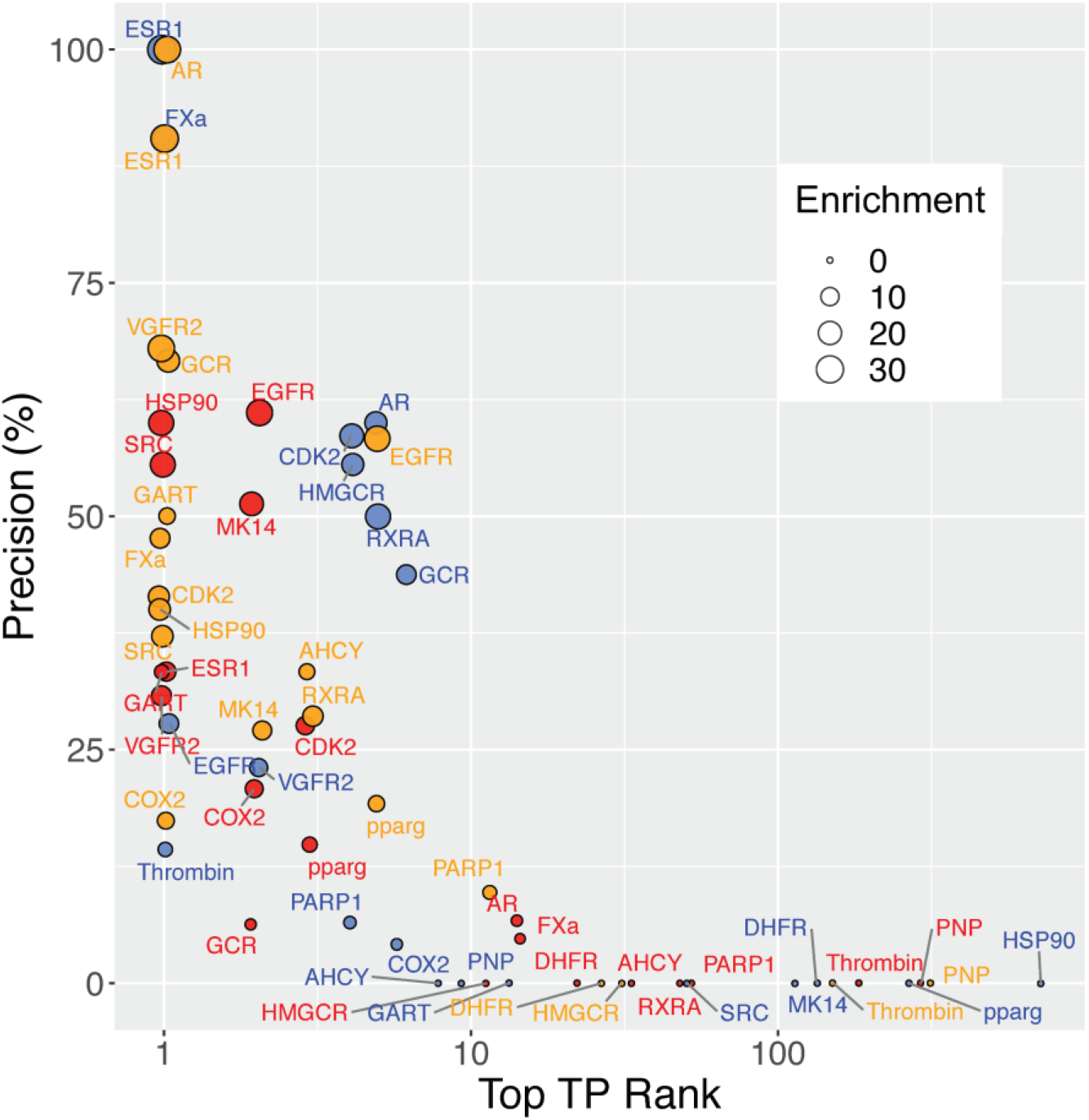
Precision at the most reduced library size (0.1%) vs. the top true positive rank for individual 20 DUD-E targets, using *Sig2Lead* (Red), *AutoDock* (Blue), and C_1_ consensus approach (Yellow) with the size of circles representing the fold enrichment at 0.1% library size.

**Figure 7:**
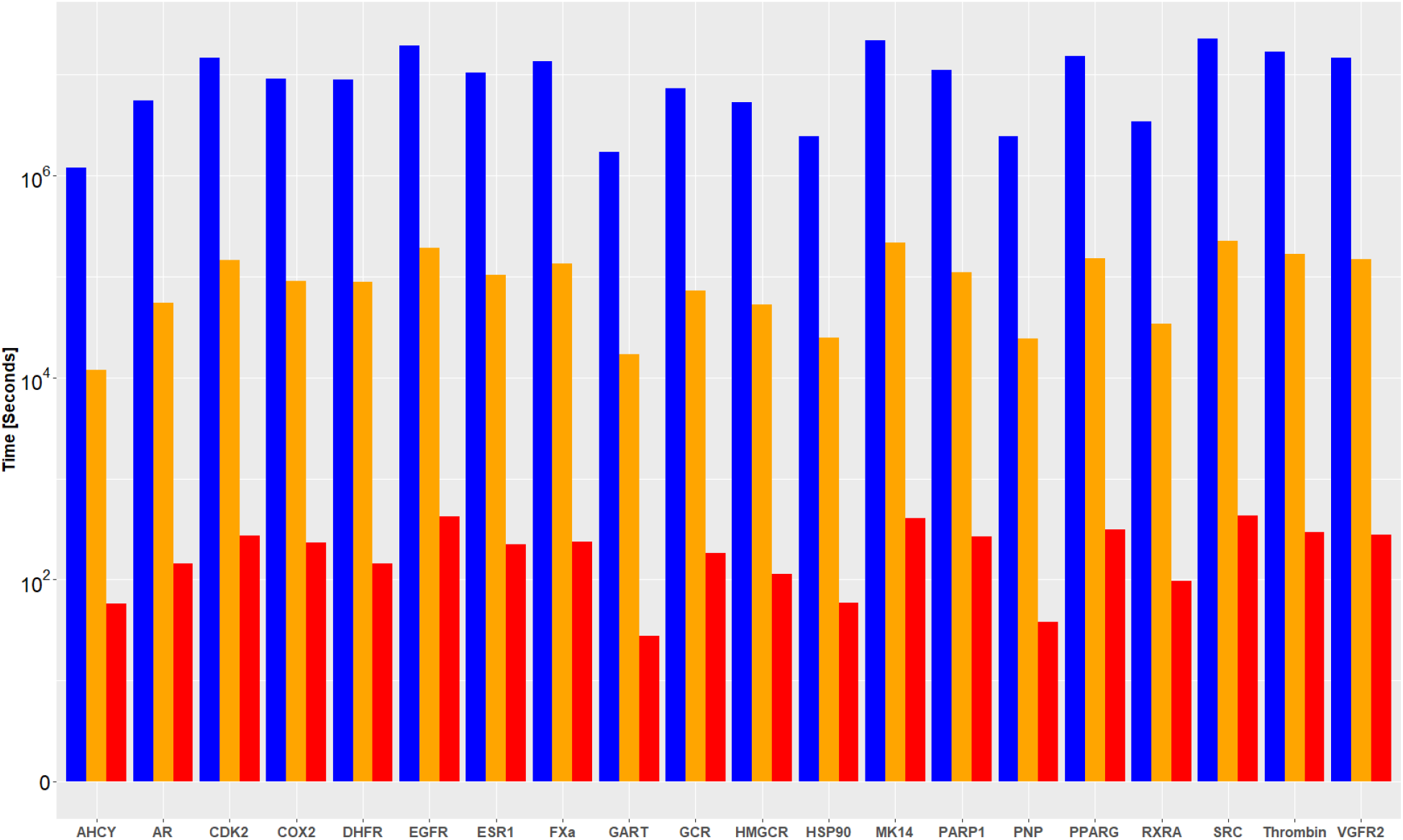
CPU times (in logarithmic scale) on 20 DUD-E targets for *ce*SAR and docking. *Autodock* alone (Blue, A) requires the longest computational time across all targets, the consensus approach (Yellow, C_1_) reduces the time by 100X, while *Sig2Lead* alone (Red, S) reduces the running by an average of 48,000X compared with docking.

It should be emphasized that the success of *ce*SAR is not due to overrepresentation of known binders from DUD-E datasets among the LINCS compounds. As can be seen from Supplemental Table 2, both true binders and decoys from DUD-E have a similar overlap with the

LINCS library, as indicated by similar distributions of Tanimoto coefficients for the closest LINCS analog for both subsets. Furthermore, the baseline approach that ignores the signal coming from signature connectivity and simply uses the Tanimoto coefficient to the closest LINCS analog, irrespective of its ‘concordance’ with the target KDs, performs consistently very poorly (see Figures 2, 4 and 5).

### Accelerating the identification of BCL2A1 inhibitors using ceSAR

Even though such an approach is less efficient, the *ce*SAR approach can be extended to incorporate signature connectivity-based re-scoring after first using docking for virtual screening to reduce the library size (this is different from the consensus approaches C considered above by reversing the order of library reduction). This form of combined approach is tested here in the context of an effort to identify specific inhibitors of an important anti-apoptotic target, namely BCL2A1 (A1). A1 has been implicated in a wide array of diseases, ranging from inflammation associated with pre-term birth^27^ to chemotherapeutic resistance in melanoma^28^. To date, no inhibitors specific to A1 have been identified and most that target the BCL2 protein family are unable to effectively block A1 activity.

Most anti-apoptotic proteins prevent apoptosis by physical binding and sequestration of pro-apoptotic proteins, achieved via binding to their “BH3” domain^29^. A major success in targeting this family was the development of a Bcl-2 inhibitor ABT-737^30^, which was modified to a bioavailable version called ABT-263 or navitoclax. Unfortunately, ABT-263 also bound Bcl-xL whose role in promoting survival of platelets, lead to thrombocytopenia in humans^31,32^. This observation spurned a biochemical tour de force that resulted in the development of ABT-199, which lost specificity for Bcl-xL^33^. Thus, despite their structural similarity, it is possible to selectively target individual Bcl-2 family members. However, few inhibitors have been developed against Bcl2-A1.

To address this challenge, a small molecule compound library of 90,087 drug-like small molecules were screened using *Autodock* v. 4.2.6. The top 300 compounds found by docking were clustered and representatives of each cluster were tested *in vitro* using a differential scanning fluorimetry thermal shift assay to detect compound binding to BCL2A1 and a fluorescence polarization competition assay to test for inhibition of Noxa BH3 domain binding to BCL2A1. Compounds were classified as inhibitors for the sake of benchmarking the *ce*SAR method if they caused a thermal shift upon addition to the BCL2A1-Noxa reaction and had an IC50, as defined by dose-response fluorescence polarization, of 400 μM or less (detailed methods available in the *Supplementary Materials*).

*Sig2Lead,* an R Shiny implementation of *ce*SAR, was then applied to re-score the tested compounds (Figure 8), demonstrating an improvement in the overall precision when ranking the top *in vitro* validated compounds. Thus, re-scoring candidate compounds obtained using docking simulations can yield further enrichment into true positives and limit the number of compounds that need to be tested experimentally. Conversely, the observed enrichment into true positives for an important and challenging target (with available LINCS KD signatures) illustrates how a set of experimentally identified weak binders can be used to seed the signature connectivity-based *ce*SAR search with the goal of identifying additional candidate compounds, i.e., the ‘concordant’ LINCS analogs of the compounds tested here.

**Figure 8:**
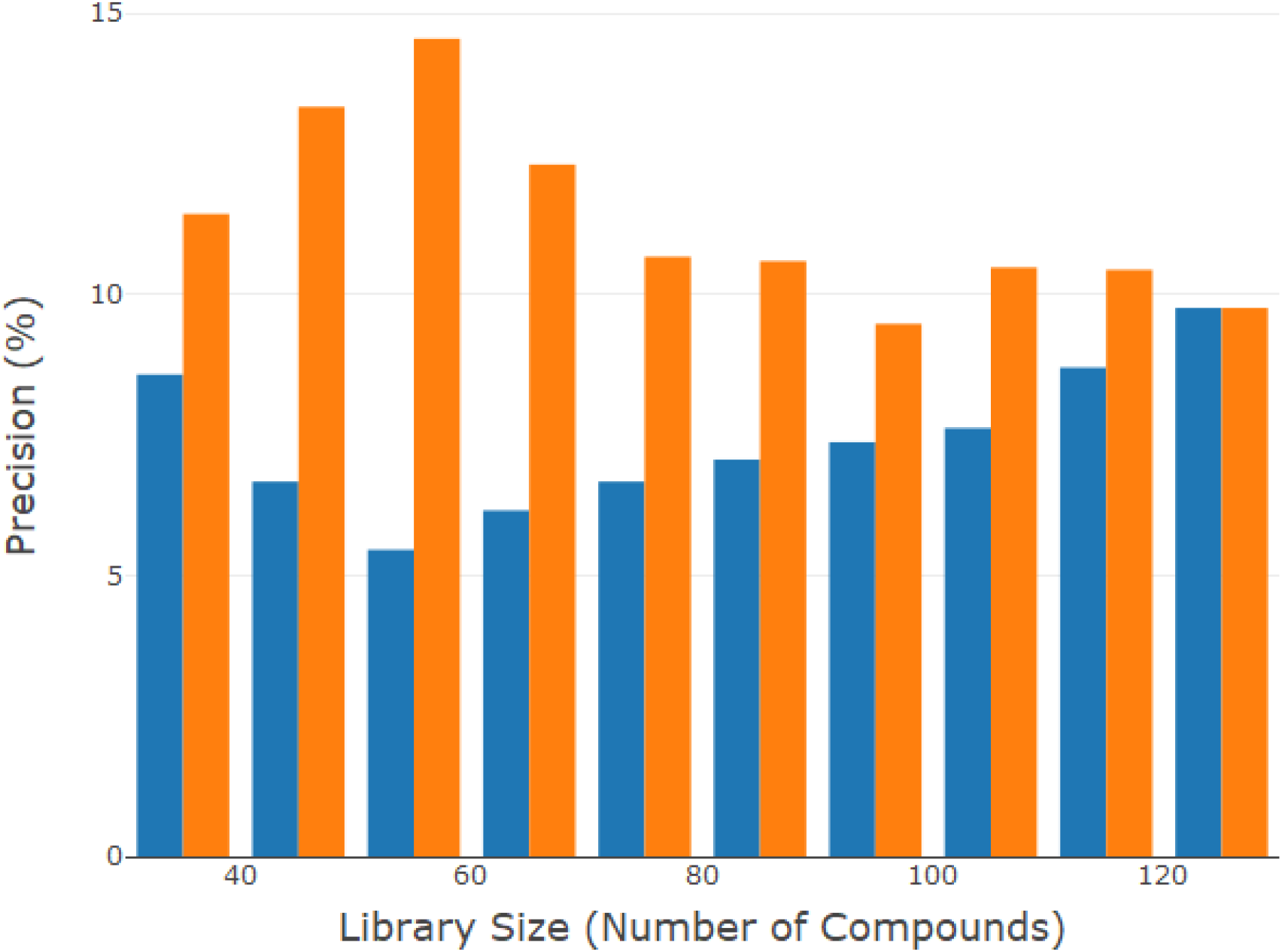
Retrospective re-scoring of a drug-discovery pipeline for BCL2A1 yields an improved enrichment into experimentally validated inhibitors using a combined *ce*SAR approach *(AutoDock* followed by *Sig2Lead* re-scoring, Yellow), as compared to docking alone (*AutoDock*, Blue).

## Discussion

Accelerating drug discovery, development and repurposing is paramount for advancing treatment options and for improving response to public health crises, such as the SARS-CoV2 pandemic, and for further progress in personalized precision medicine. *In silico* screening of small molecule libraries for their predicted interaction and inhibition of protein targets is often used to reduce the time and cost requirements in drug discovery and repurposing projects. The adage that structure dictates function has been applied in relation to small molecule inhibitors to enhance virtual screening by searching similar compounds and exploring structure activity relationships (SAR)^17,18,34,35^. In this contribution, we introduce an efficient *in silico* method to accelerate drug discovery and repurposing, dubbed *connectivity enhanced* Structure Activity Relationship (*ce*SAR). *ce*SAR improves on existing approaches by combining small molecule docking simulations with signature connectivity analysis to reduce both false positive rates and the computational cost of virtual screening, and thus allows one to overcome two major limitations of current virtual screening approaches.

Over the last two decades, transcriptional and other profiles of drug activity have been increasingly used in drug design, mode of action identification and SAR type analyses^4,12,34,36,37^. For example, identifying targets for small molecules (and thus identifying these molecules as novel inhibitors) can be facilitated by comparing bioactivity profiles or transcriptional signatures of a compound to known inhibitors^34,38^. Another important example is the use of the connectivity map approach to connect gene expression profiles of disease states (such as drug resistant forms of cancer) with discordant drug signatures, allowing one to identify drugs that can potentially be used to reverse the disease signature^4,12,15^.

The critical difference compared to these previous efforts is that *ce*SAR directly connects the transcriptional signatures of a small molecule with the signature of a gene knockdown for the purpose of identifying antagonists (or overexpression to identify agonists – an option not benchmarked in this manuscript) of a specific target rather than a pathway, and to that end combines signature connectivity analysis with atomistic docking simulations to use predicted binding energies to filter out likely pathway inhibitors. For the signature connectivity analysis, *ce*SAR capitalizes on the LINCS library of signatures which is at present the most comprehensive big data resource for pharmacogenomics^12,13^. The central advantage of LINCS is that it made available for the first time a large library of both chemical (small molecule) perturbation signatures as well as genetic (KD) perturbation signatures in one or multiple biological contexts (cell lines).

Furthermore, *ce*SAR integrates chemical similarity and signature connectivity analyses to increase the overall success rates and expand virtual screening and SAR analysis to other suitable libraries of compounds, including those identified in high throughput experimental screening. Toward that end, an algorithm for fast (exact) chemical similarity search and database retrieval is introduced that optimally exploits the sparse representation of chemical moieties as binary fingerprints. Docking and docking-based *in silico* screening, on the other hand, rely on shape complementarity between putative inhibitors and target proteins, requiring structural information about the proposed target and its relevant conformational states which may be unavailable^39,40^. The results of docking simulations may also be very sensitive to the choice of a target’s conformation, choice of the empirical force fields, docking programs and sampling depth^4,12,15^. Additionally, traditional virtual screening approaches are computationally expensive and require significant computing resources.

For example, the average CPU time per DUD-E target required to perform *AutoDock* benchmarking (with the search depth and grid sizes defined in the *Online Methods*) was of the order of 3,000 CPU hours on a computational cluster consisting mostly of 16 Intel(R) Xeon(R) CPU E5-2667 v3 @ 3.20 GHz core nodes. For comparison, the average CPU time per target required for the simple *ce*SAR search (S) was about 3.7 CPU minutes on a laptop computer with two Intel i5-4200U @ 1.6 GHz cores, and thus was reduced by roughly 50 thousand-fold compared to docking (see Figure 7). This dramatic speed increase makes it possible to perform *in silico* enrichment on large chemical libraries using a personal computer within minutes compared to weeks on a computing cluster, democratizing further the search for new drugs.

Despite its negligible computational cost, the simple *ce*SAR search (S) outperforms docking for 9 out of 20 DUD-E targets and achieves the same median precision of about 10% at extreme library reductions. The C_1_ consensus based method yields higher precision and enrichment into true binders compared to docking alone in 12 out of 20, and performs on par with docking on 3 other DUD-E targets, while still greatly reducing the overall computational cost compared to docking alone. In addition, C_1_ performs better or on par with docking for several targets (AHCY, AR, FXa) on which the simple *ce*SAR search (S) performs poorly. Importantly, the consensus approach is also more robust, as indicated by the failure rate at the extreme library reduction, defined here as the fraction of targets for which the precision is reduced to 0% at 0.1% library size. As illustrated in Figure 6, such defined failure rate is 40% (8 out of 20 targets) for *AutoDock* (A), 35% (7 out of 20 targets) for the simple signature connectivity approach (S), and 20% (4 out of 20 targets) for the consensus approach (C_1_). Another measure of failure is the number of targets for which none of the true positives is ranked among the top 100 candidates, which is 6 for *AutoDock* as opposed to 2 for *Sig2Lead* and 4 for C_1_ (it is worth noting that this number is zero for other consensus approaches – see Figure 3). Taken together, these results strongly indicate the complementarity of signature connectivity and docking based approaches for drug discovery.

On the other hand, *Autodock* (A) clearly outperforms signature connectivity enhanced methods (S and C) in 3 cases: HMGCR, Thrombin and PNP, none of which have close analogs in LINCS of the true binders included in the respective DUD-E datasets (see *Supplemental Table 2)* and/or are characterized by weak concordance between LINCS small molecule and KD signatures, which can be used to predict the likelihood of success of *ce*SAR (see *Supplemental Figures 4 and 5*). This underscores one of the obvious limitations of *ce*SAR. Namely, in addition to target gene knock-down signatures, *ce*SAR requires a representative set of transcriptionally profiled molecules broadly covering the drug-like universe. While this is largely true about the LINCS library of over 40,000 compounds (of which some 25,000 have been profiled to date), not all classes of drugs are well represented, or may not induce sufficiently strong signatures to be considered for the connectivity analysis. On the other hand, some classes of targets and their antagonists, including kinase inhibitors, are well represented in LINCS, contributing to the high accuracy of *ce*SAR on the 5 kinases included in our evaluation. For these kinases, *ce*SAR (both S and C_1_) yield improvements over docking already at 5% library size, and for the most reduced library size, achieve about 2-fold increase in median precision, which is about 50% for *Sig2Lead* alone compared to about 25% for *AutoDock* (see *Supplemental Figure 8*).

Using *ce*SAR, through the integration of signature connectivity analysis, fast exact chemical similarity search for sparse binary fingerprints, and virtual screening approaches, a dramatic increase in speed is obtained while improving accuracy, thus providing a fast, robust and accurate platform for drug discovery and repurposing. We believe that the performance of *ce*SAR adds significantly to the utility of LINCS as a big data resource for pharmacogenomics and provides a strong argument in favor of further large-scale transcriptional profiling of drug-like molecules and druggable parts of the genome. We anticipate that with further advances in the CRISPR technology, more accurate gene signatures will be obtained, leading to increased performance of the new approach. At the same time, continued advances in determining 3D structures of proteins and their complexes by using cryo-electron microscopy and other techniques will expand the protein targetable space, adding to the importance of accelerating the speed of virtual screening approaches.

## Methods

### Candidate molecule ranking using ceSAR

*ce*SAR ranks candidate molecules by combining signature connectivity analysis and chemical similarity search to identify the most similar ‘concordant’ LINCS analogs of candidate compounds. Here, ‘concordant’ is defined as having a signature that is significantly positively correlated with a target gene knock-down signature. For a target gene *t*, with at least one knock-down transcriptional signature available in LINCS, *t ∈ L*, and for a library of small molecules to be ranked, *Q*, the following similarity score is computed for each *q* ∈ *Q* as a basis for ranking:

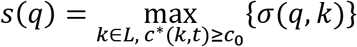

where *σ*(*q, k*) is the Tanimoto coefficient (Jaccard similarity measure)^19^ between compounds *q* and fc ∈ *L* represented as binary fingerprints, while *c*\*(*k, t*) is the maximum concordance (over all cell lines for *t*, and cell line, concentration, exposure time tuples for *k*) between the signatures of chemical perturbagen fc and genetic knock-downs of *t*:

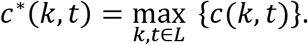

Conceptually, taking the maximum value of signature concordance over all cell lines and concentrations (for chemical perturbagens) follows the assumption that genetic and chemically induced loss of function may result in the most pronounced signatures and concordance in some unknown biological contexts, as represented by different cell lines included in LINCS. Pearson correlation coefficient (PCC)-based concordance measure, *c*(*k, t*) = *PCC*(*k, t*), is used here^23^, and the threshold for significant concordance is set to *c*_0_ = 0.2. As shown in *Supplementary Materials,* the performance of the method is robust with respect to the choice of this threshold (see, e.g., *Supplemental Figure 9*).

### Sparse binary fingerprints for chemical similarity search

Binary fingerprints are widely used in cheminformatics for efficient chemical similarity search and SAR analyses^41–44^. In this approximation, small molecules are represented as binary vectors indicating the presence of substructures, subgraphs, pharmacophores or chemical groups^41,42^. Here, we use the 1024-bit atom-pair fingerprint representation^41,45^, as generated by the *ChemmineR* package^46,47^, which leads to a sparse binary vector representation of LINCS compounds. Indeed, as shown in *Supplemental Figure 1*, very few of the fingerprint features have a relatively balanced split between ones and zeros across the LINCS compounds. In addition, all LINCS compounds have less than 120 ones in their respective fingerprints of length 1024, with median of about 50 ones.

### Fast exact chemical similarity search using minSim

Consider now a search for a query compound *q* ∈ *Q* against a database compounds *k* ∈ *L* using binary fingerprints described above. The formula for the Tanimoto coefficient, *σ*(*q, k*), which is defined for two binary fingerprints *q* and *k* as the ratio of the number of positions with ones in both *q* and *k* and the number of positions with ones in either *q* or *k,* can be written in the following form:

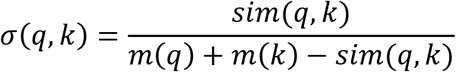

where *m*(*q*) and *m*(*k*) are the number of ones that can be pre-computed for all database molecules *k*, while *sim*(*q, k*) is the number of ones in common for *q* and *k*.

Note that the computation of *sim*(*q, k*) can be limited to only those columns in the binary fingerprint where *q* is in the minority state, which is assumed to be 1. Furthermore, by using preprocessing of the reference data set of compounds (here LINCS library) one can optimally exploit the sparsity in each column by pre-computing indexes of database compounds in the minority state at each column, as illustrated in *Supplemental Figure 2*. Namely, the following list of database vectors *k_i_* is pre-computed for each column *j* in the fingerprint:

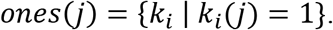

The *minSim* (for *minority Sim)* algorithm, computes all Tanimoto coefficients for a query molecule *q* by updating integer counters *sim*(*q, k*), which are set to zero for all *k* at the beginning of the search, in a simple loop over minority columns in *q* and minority lists in each minority column:

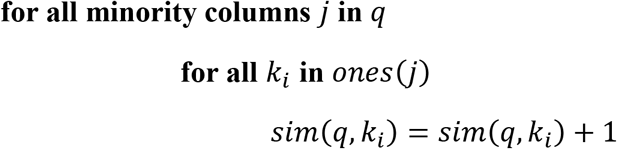

We posit that *minSim* optimally exploits the sparse nature of binary fingerprints by considering only those fingerprint columns (positions) where the query molecule *q* is in the minority state, and by using precomputed lists of all database compounds fc that are in the minority states at these positions. The implementation of the algorithm in R is included in *Supplemental Figure S3.*

Note also that *minSim* computes the exact Jaccard similarity, without using approximate techniques, such as those based on hashing^25,26,46^. As can be seen from Supplemental Table 1, for the retrieval from the LINCS library for different DUD-E datasets, *minSim* provides between 60 and 150-fold speed-up compared to fpSim function, which represents traditional approaches for exact chemical similarity search^46^. Note that these speed-ups are consistent with the observed levels of sparsity in the LINCS dataset, while reflecting the varying degree of sparsity in DUD-E datasets of query molecules.

### Statistical analysis

Multiple comparison was performed by Kruskal-Wallis test, while the median difference of the precision between two methods was assessed by the two-sided Wilcoxon test. The difference of the whole distribution of the precision values at different sizes of the reduced libraries between methods was performed by using the two-sided Kolmogorov-Smirnov test.

## Data Availability

DUD-E data sets and target structures used here for benchmarking can be downloaded from http://dude.docking.org/subsets/dud38. The LINCS library of small molecules can be downloaded from the LINCS Data Portal (http://lincsportal.ccs.miami.edu/dcic-portal/) while its pre-processed counterpart for fast chemical similarity search and SAR analyses can be downloaded from https://github.com/sig2lead. The gene knock-down and chemical perturbation LINCS signatures, as well as their pre-computed concordance scores are available through *iLINCS* and its API programmatic interfaces (http://www.ilincs.org).

## Code Availability

*ce*SAR has been implemented as an *R Shiny* app, dubbed *Sig2Lead,* that uses API calls to *iLINCS* to obtain concordance scores for LINCS compounds and a target gene, and precomputed indexes for fast retrieval of LINCS analogs using the *minSim* algorithm. *Sig2Lead* is a public domain package and can be downloaded from https://github.com/sig2lead.

## Acknowledgements

This work was supported in part by the National Institutes of Health grants U54 HL127624, P30 ES006096, R01 MH107487, R01CA122346, R01GM128216, 1T32CA236764, R01 CA237016, R21 HD090856 and UL1TR001425, 2I01BX001110 BLR&D VA Merit award, and Cincinnati Children’s Innovation Fund award (to DAH and ABH).

## Author Contributions

AT, JR and SC developed and benchmarked the new method, contributed several sections of the manuscript and implemented the *Sig2Lead* app, JR, RA and BS developed and tested the *minSim* algorithm, BS, MP, MFN, MK and MM developed the *i*LINCS platform and API interfaces for signature connectivity analysis used by *Sig2Lead*, AT, AH and DH contributed the BCL2A1 experimental validation, AM, MCK, RM, WS, NS, NN, YZ helped with the benchmarking and refinement of the method and the manuscript, JM conceived of the method and coordinated the study and writing of the manuscript.

